# Convergent evolution of niche structure in Northeast Pacific kelp forests

**DOI:** 10.1101/2020.02.24.963421

**Authors:** Samuel Starko, Kyle W. Demes, Christopher J. Neufeld, Patrick T. Martone

## Abstract

1. Much of the morphological and ecological diversity present on earth is believed to have arisen through the process of adaptive radiation. Yet, this is seemingly at odds with substantial evidence that niches tend to be similar among closely related species (i.e., niche conservatism). Identifying the relative importance of these opposing processes in different circumstances is therefore essential to our understanding of the interaction between ecological and evolutionary phenomena.
2. In this study, we make use of recent advances in our understanding of the phylogeny of kelps (Laminariales) to investigate niche evolution in one of the most important groups of benthic habitat-forming organisms on the planet. We quantify functional traits and use community sampling data from a kelp diversity hotspot to determine which traits are responsible for the habitat (β) niche of kelps and whether they are labile or conserved across the kelp phylogeny.
3. We find that combinations of functional traits have evolved convergently across kelp subclades and that these traits are significant predictors of community structure. Specifically, traits associated with whole-kelp structural reinforcement and material properties were found to be significantly correlated with species distributions along a gradient of wave disturbance and thus predict the outcome of environmental filtering. However, kelp assemblages were made up of species that are more phylogenetically distinct than predicted from null models (i.e., phylogenetic overdispersion), suggesting that niche partitioning along this gradient of wave disturbance has been an important driver of divergence between close relatives.
4. These results collectively demonstrate that environmental filtering by waves plays an essential role in determining the habitat niche of kelps across local communities and further suggest that this community-level process can drive phenotypic divergence between close relatives. We propose that parallel adaptive radiation of kelp subclades has shaped the diversity and species composition of kelp forests in the Northeast Pacific and we discuss how evidence from the literature on incipient or ongoing speciation events support this hypothesis.

## Introduction

A major challenge among ecologists is to understand how community-level processes influence the macroevolution of lineages (Webb et al. 2002, Emerson and Gillespie 2008, Gerhold et al. 2015). Local environmental gradients serve as the environmental context in which both ecological and evolutionary processes occur and can thus serve as a starting point to address this challenge. In the context of communities, stress and/or disturbance from the environment can exceed the tolerances of some species, causing them to be excluded from certain communities (e.g. van der Valk 1981, Menge and Sutherland 1987, Webb et al. 2002, Cornwell and Ackerly 2009, Kraft et al. 2014). Thus environmental gradients can serve as “environmental filters”, resulting in communities of species that share phenotypic traits necessary to survive in a particular environment (Reich and Oleksyn 2004, Swenson and Enquist 2007, Kraft et al. 2011, 2014, Enquist et al. 2015, Cavalheri et al. 2015, Ulrich et al. 2017). Over evolutionary timescales, environmental gradients can influence the phenotypic evolution of community members by serving as strong sources of selective pressure (Cavender-Bares et al. 2004a, Demes et al. 2013, Gerhold et al. 2015). Thus, community assembly dynamics along environmental gradients depend strongly on the interplay of these ecological and evolutionary processes. Yet, disentangling the factors at play has been an ongoing challenge (Cavender-Bares et al. 2009).

Depending on the evolutionary history of the species pool and the evolutionary lability of underlying phenotypes, we might expect very different patterns of relatedness among the species found in local communities subject to environmental filtering. Many studies have found that closely related species share similar phenotypes (Webb 2000, Webb et al. 2002, Silvertown et al. 2006a, Kraft et al. 2007) due to selection against phenotypic divergence (“niche conservatism”) or due to a lag caused by a shared ancestor and slowly evolving traits (Wiens 2008, Losos 2008). This pattern is remarkably common (Darwin 1859, Webb et al. 2002, Vamosi et al. 2009), leading many researchers to assume that it is true, even in the absence of any phenotypic data (see Gerhold et al. 2015 for a review). When phenotype and phylogeny are correlated, closely related species are often clustered in space because close relatives with similar traits tend to experience similar outcomes from strong environmental filtering (Fig 1a; Webb et al. 2002, Cavender-Bares et al. 2009). However, the ubiquity of studies showing evidence for niche conservatism stands in contrast to another body of work on the process of adaptive radiations wherein lineages are known to spread out across environmental gradients (hereafter “niche partitioning”) to move into open niches as they diversify (MacArthur 1958, Hector and Hooper 2002). This process would be expected to result in the opposite community pattern: communities made up of distantly related species that share a set of convergently evolved traits (Fig 1b; Cavender-Bares et al. 2004a, Silvertown et al. 2006a, 2006b, Cavender-Bares et al. 2018). In order to reconcile this apparent disconnect between alternative theoretical expectations of phylogenetic community structure, it is necessary to determine the relative importance of these opposing evolutionary forces (niche conservatism versus niche partitioning) in various lineages and circumstances to determine how and when particular processes dominate phenotypic evolution. The relative importance of these different processes can be inferred by identifying the patterns of phenotypic variation across the phylogeny of a given lineage and by determining how this phenotypic variation relates to the sorting of species into ecological communities (Lopez et al. 2016).

**Fig 1.**
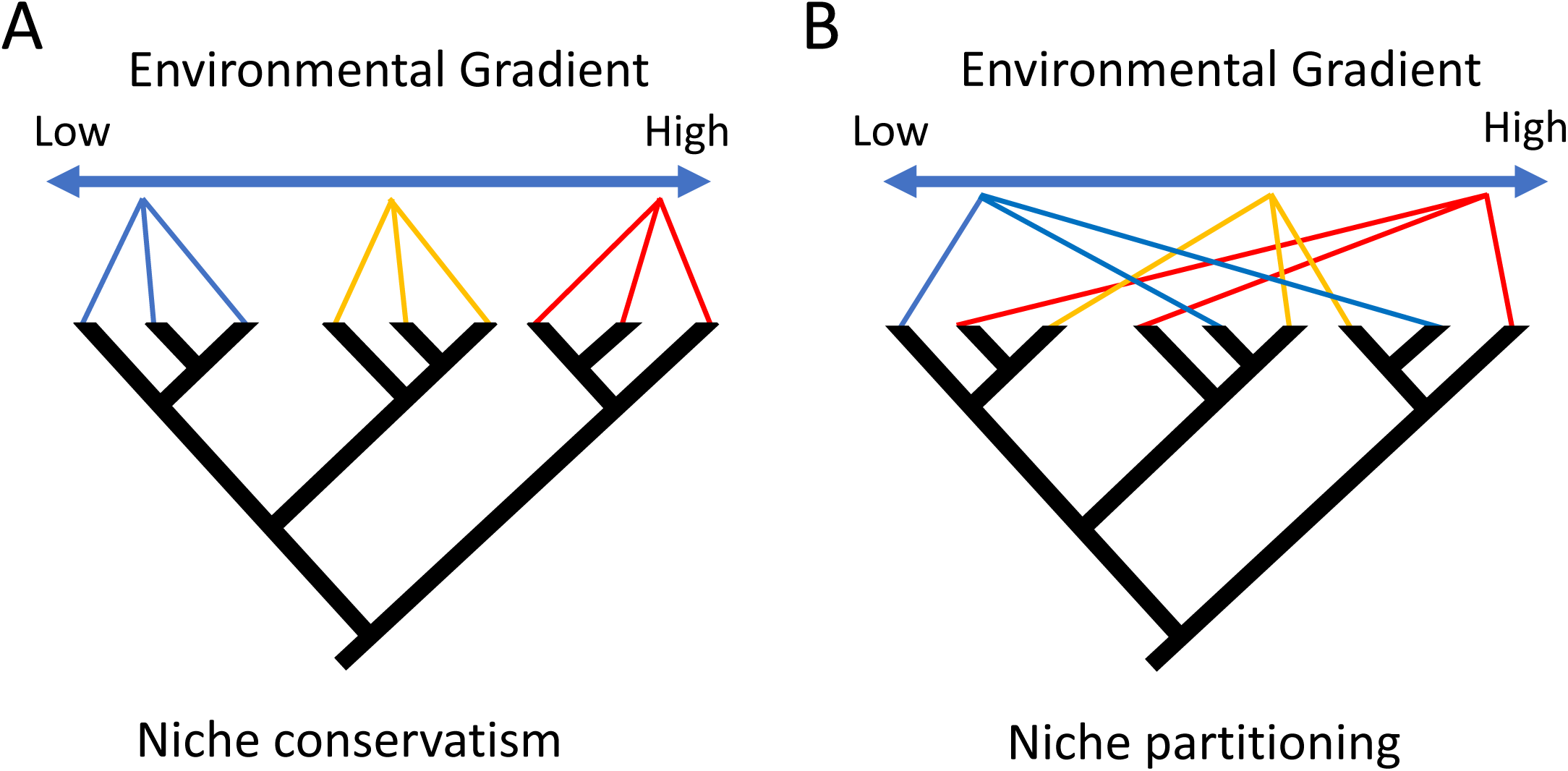
Theoretical extremes of how communities might be phylogenetically structured along environmental gradients under different dominant evolutionary processes. Lines are drawn from tips of the phylogeny to one of three communities situated along a theoretical disturbance gradient. Colours indicate a particular set of traits and environmental filtering drives trait clustering in both examples. If niches are conserved within subclades, then communities are expected to be clustered phylogenetically (Panel A; e.g., Webb 2000). If close relatives partition niches across the environmental gradient, then communities are expected to be phylogenetically overdispersed (Panel B; e.g., Cavender-Bares et al. 2004a).

While the relatedness of species within and between communities (hereafter, phylogenetic community structure) has been well explored in terrestrial taxa, particularly embryophytes (Emerson and Gillespie 2008, Cavender-Bares et al. 2009), most marine lineages are poorly studied in this respect (Verbruggen et al. 2009, Best and Stachowicz 2013). This is problematic because evolutionary processes in the ocean may be somewhat different from those on land, with generally fewer barriers to reproduction in marine environments (Buzas and Culver 1991, Schluter 2000). Marine macroalgae offer an intriguing study system to explore the evolution of phenotype and niche structure because morphologies, which are relatively simple, strongly influence the abiotic tolerances of species (Littler and Littler 1984, Steneck and Dethier 1994, Martone 2007, Starko and Martone 2016). In particular, water motion from waves and currents is believed to act as an exceptionally strong environmental filter that excludes species from more wave exposed sites if they are not strong enough to resist the forces that they experience (Denny 1985, Gaylord et al. 1994, Denny and Gaylord 2002, Demes et al. 2013). Conversely, low flow habitats may be highly stressful due to the formation of diffusive boundary layers that reduce nutrient uptake and gas exchange across macroalgal thalli (Hurd 2017). Thus, low flow environments may eliminate species that fail to achieve morphologies that facilitate the depletion of boundary layers when water motion is low (Coyer and Roberson 2004). This continuum of stress and disturbance caused by the position of local communities along gradients of water motion is an essential driver of both community assembly processes and the evolution of phenotypic traits across rocky shores, but ecological and evolutionary processes have yet to be linked across any major lineage that occupies this environment.

Kelps (order Laminariales) are the largest and most productive macroalgae in the ocean and dominate approximately 25% of coastlines globally (Wernberg et al. 2019). Kelps increase the productivity of cool, temperate nearshore ecosystems and their presence can substantially alter the composition of biotic communities (Steneck et al. 2002, Graham 2004, Teagle et al. 2017, Hind et al. 2019). In spite of their global importance, we still have a limited understanding of the processes underlying the evolution of kelps. While recent advances in phylogenetics have dramatically improved our understanding of the relationships between species and the evolution of some key morphological features (e.g., Lane et al. 2006, Kawai et al. 2013, Jackson et al. 2017, Starko et al. 2019b), it is unknown how niche structure has evolved across this ecologically diverse clade. Kelps diversified in the North Pacific following the Eocene-Oligocene boundary (Starko et al. 2019b), possibly as a result of ecological opportunity that arose as the North Pacific cooled over the past 30 million years. While kelps are found globally, they are overwhelmingly most diverse in the North Pacific and it remains largely unclear what processes have allowed for the production of such high sympatric diversity in this part of the ocean.

In this study, we investigate the phylogenetic patterns of habitat (β) niche structure across geographically co-existing species of kelp in the Northeast Pacific, one of the most diverse stretches of coastline for kelps and their likely center of origin (Starko et al. 2019b). We begin by presenting a dataset of quantitative traits for 17 species of kelp and testing for phylogenetic signals on these traits. We use an ancestral state reconstruction approach to determine whether particular trait combinations share a common origin or whether they have convergently arisen in different subclades. Next, we test whether environmental filtering is an important driver of community assembly and determine how this relates to the phenotypic and phylogenetic structure of communities. We do so by making use of a community dataset that spans a gradient of wave action, an important driver of nearshore community composition and a known filter of the kelp species pool (Duggins et al. 2003, Burel et al. 2019). By teasing apart the evolution of phenotypic features from patterns of phylogenetic community structure, our results lend critical insights into the evolution of niche structure across one of the most ecologically important groups of foundations species found anywhere in the ocean and shed light on how ecological and evolutionary forces interact to shape marine communities.

## Materials & Methods

### Quantifying phenotypic traits

Seven quantitative traits were compared for all kelp species of interest (n=17), many of which are analogous to commonly measured traits in land plants; these included two traits describing whole individual biomass allocation (stipe mass fraction or SMF, holdfast mass fraction or HMF) and five traits describing mechanical and structural properties of blade tissues. SMF and HMF describe the proportion of total biomass that is stipe or holdfast material, respectively. HMF is analogous to root-shoot ratios in land plants. Organs (holdfast, blades, stipes) of individual kelps (n = 5 per species) were carefully separated and dried in a 50-60°C drying oven. Blade mass per area (BMA; analogous to leaf mass per area) was defined as the amount of dry biomass per unit area of blade tissue and dry matter content (DMC) was defined as the ratio of dry weight to wet weight. Both BMA and DMC were measured by taking hole punches of standardized area out of the blades and measuring the wet mass and dry mass of each hole punch. Mechanical properties of blade material: breaking stress (σ), stiffness (*E*) and extensibility (ε), were measured using an Instron (model 5500R, Instron Corp., Canton, Massachusetts, USA), a portable tensometer (described in Martone 2006), or were taken from the literature (Tables S1-S2). With the exception of these few material properties measurements taken from the literature, trait data represent average measurements taken from adult individuals of populations in southern British Columbia (Barkley Sound, Port Renfrew, Vancouver or Victoria; see Tables S1-S2). We used a principal components analysis to collapse trait combinations into fewer axes of correlated traits. Then, to determine whether any major PCA axis correlates with the ability of kelps to resist dislodgement, we tested for correlations, using PGLS models, between PCA axes and tenacity-area scaling relationships quantified previously (Starko and Martone 2016) for the 8 species included in that study. Tenacity-area scaling relationships describe the slope of the relationship between maximum dislodgement force and thallus size and are therefore an effective measure of wave tolerance.

**Table 1.** Statistical testing of phylogenetic signal for quantitative traits

| Functional Traits | Phylogenetic Signal |  |  |  |
| --- | --- | --- | --- | --- |
|  | Blomberg's K | P-value | Pagel's Lambda | P-value |
| PC1 | 0.538 | 0.610 | <0.01 | >0.99 |
| PC2 | 0.612 | 0.425 | <0.01 | >0.99 |
| HMF | 0.353 | 0.693 | <0.01 | >0.99 |
| SMF | 0.860 | 0.063* | 1.128 | 0.085* |
| BMA | 0.718 | 0.190 | <0.01 | >0.99 |
| DMC | 0.521 | 0.649 | <0.01 | >0.99 |
| Strength | 0.584 | 0.457 | 0.108 | 0.737 |
| Stiffness | 0.720 | 0.197 | 0.303 | 0.437 |
| Extensibility | 0.285 | 0.962 | <0.01 | >0.99 |

\*Trending towards significance ( $P < 0.10$ )

### Phylogenetic reconstruction

The phylogeny of kelps, with more than 120 species, has been studied previously in considerable detail (Lane et al. 2006, Jackson et al. 2017, Starko et al. 2019b). In this study, the time-calibrated phylogeny inferred by Starko et al. (2019b) was used to represent phylogenetic divergence in millions of years for the 17 co-occurring Northeast Pacific kelp species of interest. This time-calibrated phylogenomic analysis is the most well supported and comprehensive to date and included all 17 species except *Laminaria setchellii*, which was incorporated into the analysis by substituting it for *L. digitata*, which is not found in the northeast Pacific but was included in the phylogenomic analysis. This substitution relies on the assumption that *L. setchellii* has an equivalent divergence time from *Laminaria ephemera* as *L. digitata*, which is well supported by previous work on intrageneric relationships between *Laminaria* species, showing less than 1 million years difference in divergence time between *L. ephemera* and *L. setchellii* vs. *L. digitata* (Rothman et al. 2017). Phylogenetic signal of traits was measured using Blomberg’s K (Blomberg et al. 2003) and Pagel’s lambda (Pagel 1999). We also tested for correlations between trait distance and phylogenetic distance using Mantel tests.

We used the software “StableTraits” (Elliot and Mooers 2014) to reconstruct ancestral values of principal component axes and the traits and to model rates of phenotypic evolution. “StableTraits” samples from a heavy-tailed distribution, therefore allowing for modelling of traits under selection. We ran StableTraits for 10 million generations, sampling every 1000 generations. Results of these analyses were visualized using the contMap function in “phytools” (Revell 2012).

### Community dataset

To determine how trait or phylogenetic differences influence community assembly, we used a community dataset of intertidal kelp distributions in Barkley Sound, British Columbia that was published in a Parks Canada technical report (Druehl and Elliot 1996). Data from sites sampled in 1995 (n = 87 sites), the most extensive year of this survey, were combined into a data matrix. This dataset included all of the species examined in the trait analysis except two (*Laminaria ephemera* and *Cymathaere triplicata*). Although a coarse categorical abundance measurement is given in their report, only presence and absence data were used. At a subset of sites (n = 55) that could be located by photographs in the 1996 report, the upper limit of barnacles was measured in the summers of 2018-2019 and these values were used as a continuous proxy for wave exposure. The upper limit of barnacles is an effective proxy of wave run-up and is known to increase in elevation at more wave exposed sites (Harley and Helmuth 2003, Neufeld et al. 2017). The upper limit of barnacles was measured by using a stadia rod and sight level, along with tide predictions from Bamfield Inlet, Effingham Island or Mutine Point, depending on proximity. A categorical measure of wave exposure provided by Druehl & Elliot was used for analyses of all 87 sites. Barnacle upper limit was significantly different between these wave exposure categories (ANOVA: F_2,52_ = 19.5815, P < 0.0001) with significant differences between all means (Tukey HSD < 0.05), suggesting that barnacle upper limits are an appropriate proxy for wave exposure. Using the range of barnacle upper elevation data (that spanned approximately 3 to 5.5 m above MLLWLT), we created a “wave exposure index” by subtracting 3 meters from each measurement and then dividing by 2.5 (the approximate range of barnacle upper limits), resulting in an index that varied from 0 to 1. Although resurveys were conducted at some of these sites, recent work demonstrated that kelp forests have been lost from several of these sites, likely as a result of the 2014-2016 heatwave (Starko et al. 2019a). Thus, only historical data were used to reconstruct niche structure before the large-scale degradation of these kelp communities.

### Quantifying species co-occurrence

First, to determine whether non-neutral processes were required to explain the distribution of species across communities, we tested whether our community matrix was significantly different from randomly generated communities. We did so by comparing our observed checkerboard score (i.e., c-score; Stone and Roberts 1990), a measure of association between species pairs, to randomly simulated communities. In order to test for significant associations between individual species, observed co-occurrence probabilities were calculated for each pair of species and compared to a null expectation of species co-occurrence that was generated using randomizations that considered only the number of sites at which each species was found. In cases where species were expected to co-occur at less than one site, these species pairs were excluded due to insufficient data. Deviations from expectations were measured using a log response ratio of observed vs. expected outcomes, hereafter “co-occurrence index”. Calculated as:

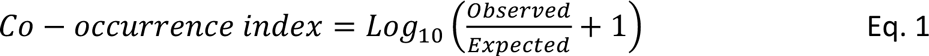

where “Observed” refers to the actual number of co-occurrences in the community matrix, and “Expected” refers to the number of sites that species were expected to be found together given the null model. Species association analyses were corrected for false detection rate and were considered significant when q-values were less than 0.05. In order to determine whether phylogenetic distance or trait differences (first and second trait-derived principal components) influenced the co-occurrence probability of species, linear regressions were fit between each predictor (phylogenetic distance, PC1 distance and PC2 distance) and co-occurrence index.

### Wave exposure and community assembly

The relationship between species presence and wave exposure was measured in two ways using the subset of sites (n = 57) for which continuous wave exposure (barnacle upper limit) had been measured. This subset did not include any sites with *S. latissima*, which was therefore excluded from these analyses. It also included only one observation of *P. palmaeformis* at one of the most wave-exposed sites in our dataset. This species is well known to occur only on the most wave exposed shores (Nielsen et al. 2006) and so this site was deemed representative of the niche of *P. palmaeformis.* However, to better improve our estimate of average wave exposure for this species, we measured the upper limit of barnacles at two sites on the nearby outer coast (Cape Beale) that consistently have *Postelsia palmaeformis* populations. All three sites were very high exposure (upper limit of barnacles: 5.2 – 5.8 m above MLLWLT). To assess the relationship between traits and species’ habitat use, average wave exposure was measured for each species from all sites in which that species was present. A phylogenetic least squares (PGLS) regression was then used to test for an effect of principal component axes and all seven quantitative traits on average wave exposure. In order to further visualize differences in species habitat use, the probability of species presence was plotted against wave exposure (i.e., the upper limit of barnacles) as modeled using polynomial, binomial generalized linear models. This modelling approach allows for an optimal wave exposure rather than forcing saturation. This was done separately for members of the two subclades with the most species included here, the families Arthrothamnaceae and Alariaceae. To determine whether sites of different wave exposure also have different kelp communities, we conducted a PERMANOVA with the wave exposure categories described above as a predictor variable.

### Phylogenetic community structure

To further test for an effect of phylogeny on community assembly we used indices of phylogenetic community structure (Webb 2000). Net relatedness index (NRI) and nearest taxa index (NTI) measure the extent to which taxa are phylogenetically clustered at a particular site relative to the regional species pool. A positive value of either NRI or NTI indicates phylogenetic clustering, while negative values indicate phylogenetic overdispersion. NRI measures phylogenetic clustering by considering the average phylogenetic distance between all members of a community. Specifically, NRI is defined as follows:

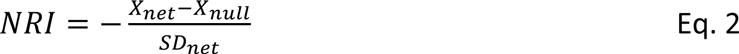

Where X_net_ is the average phylogenetic distance between members of a community, and X_null_ and SD_null_ represent the mean and standard deviation, respectively, of simulated random draws from the species pool. We calculated these metrics using 10,000 random simulation. NTI is similar to NRI but considers the average distance between each species and its closest relative. Specifically, X_net_ from equation 2 is replaced with X_min_ which is defined as the average distance between each species and its closest relative, such that:

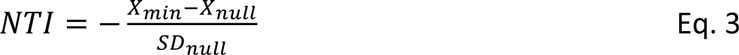

For NTI, X_null_ and SD_null_ represent the mean X_min_ and associated standard deviation from random draws of the species pool, similar to calculations of NRI. As a consequence of differences in the underlying metric of interest (X_net_ versus X_min_), NRI is more sensitive to phylogenetic clustering deeper into the phylogeny, while NRI is more sensitive to clustering near the tips of the phylogeny. The significance of trends in phylogenetic structure was evaluated in two ways. First, at a community level, sites (i.e. individual communities) were considered to be significantly structured by phylogeny if NRI or NTI values ranked among the 500 most extreme values (97.5th or 2.5th percentiles) of the 10,000 randomly generated pseudo-communities. A second approach was used to determine if, across the whole dataset, there were significant trends in phylogenetic community structure. NRI and NTI are both expected to be approximately normally distributed with a mean of zero, therefore in order to determine whether the mean of the distribution of kelp communities differed from this null expectation, t-tests were also performed, treating sites as replicates (as in Cooper et al. 2008).

### Statistical software

All statistical analyses were performed in R version 3.6.0, using the packages “ape” (Paradis et al. 2004), “phytools” (Revell 2012), “picante” (Kembel et al. 2010), “qvalue” (Bass et al. 2018), “EcoSimR” (Gotelli et al. 2015), and “cooccur” (Griffith et al. 2016).

## Results

### Phenotypic traits are convergent across taxa

Principal component analysis resulted in seven component axes with the first two explaining 63.9% of the variation in trait values (Fig 2A). Principal component 1 (PC1) correlated with structural characteristics of the whole kelp (HMF and SMF), as well as the blade (DMC, BMA), which were themselves all positively correlated (Fig S1). Principal component 2 explained mainly the properties of materials (σ, *E* and ε). These two components explained 35.3% and 28.6% of the total variation in functional traits respectively. Principal component 1 strongly correlated with tenacity-area scaling relationships (Fig S2; PGLS model: F = 11.92, df = 1 and 6, P = 0.0136). There was no significant phylogenetic signal on any of the traits investigated in this study, including principal components (Table 1, Fig 2B). However, our analysis revealed a possible but not significant phylogenetic signal on SMF (Blomberg K: 0.860, P = 0.063; Pagel’s Lambda = 1.128, P = 0.085). Although not significant, Blomberg’s K was < 1 in all cases, indicating that traits tended to be more dissimilar among close relatives than predicted from a Brownian motion model. Some pairs of closely related species were somewhat similar in at least some traits (e.g. *Pleurophycus gardneri* and *Pterygophora californica*), but for the most part, closely related species differed as much or more than distantly related ones (Fig 2B). This observation was confirmed by the lack of a significant relationship between PC1 and PC2 trait distances and phylogenetic distance (PC1 Mantel test: Z-stat = 6450.835, p = 0.589; PC2 Mantel test: Z-stat = 6449.193, p = 0.691). Ancestral state reconstruction demonstrates that trait combinations have evolved repeatedly across the kelps with clear patterns of phenotypic convergence (Fig S3).

**Fig 2.**
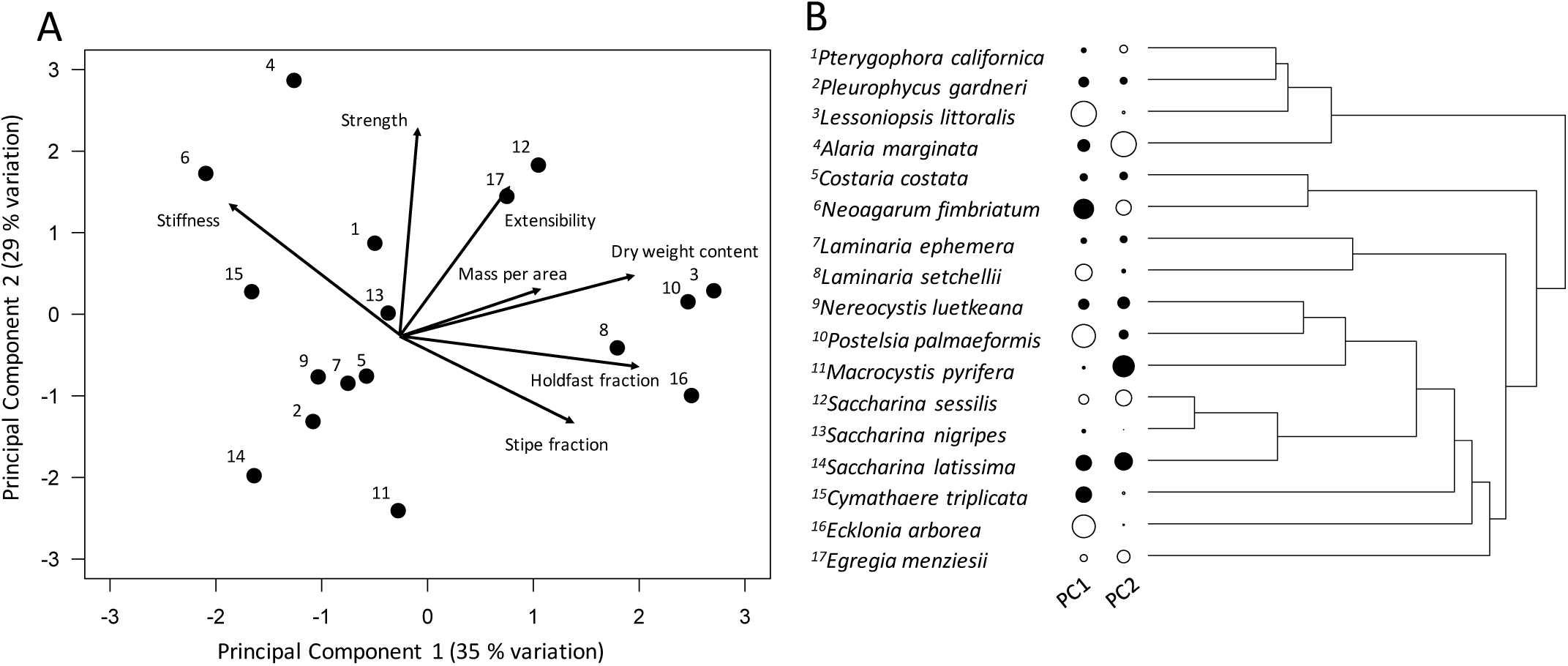
Phylogenetic distribution of trait axes in northeast Pacific kelp species. Panel A shows the first two principal component axes. Panel B shows PC1 and PC2 plotted on the phylogeny. The size of each bubble indicates the value of each trait axis and the colour indicates whether values are positive (white) or negative (black). There is no significant phylogenentic signal in either axis.

### Kelp communities are phenotypically (not phylogenetically) clustered

The community matrix was non-random with a c-score that exceeded the range of values from random simulations (Fig S4). There were several significant associations between species (Fig 3). Positive and negative species associations occurred between both closely and distantly related species pairs. For example, closely related species *Macrocystis pyrifera* and *Nereocystis luetkeana* were negatively associated with each other, while sister taxa, *Pleurophycus gardneri* and *Pterygophora californica*, were positively associated (Fig. 3). Moreover, *Egregia*, the most phylogenetically distinct genus from the family Arthrothamnaceae, was positively associated with some members of three other families (Alariaceae, Agaraceae, Laminariaceae) and negatively associated with a member of one (Agaraceae).

**Fig 3.**
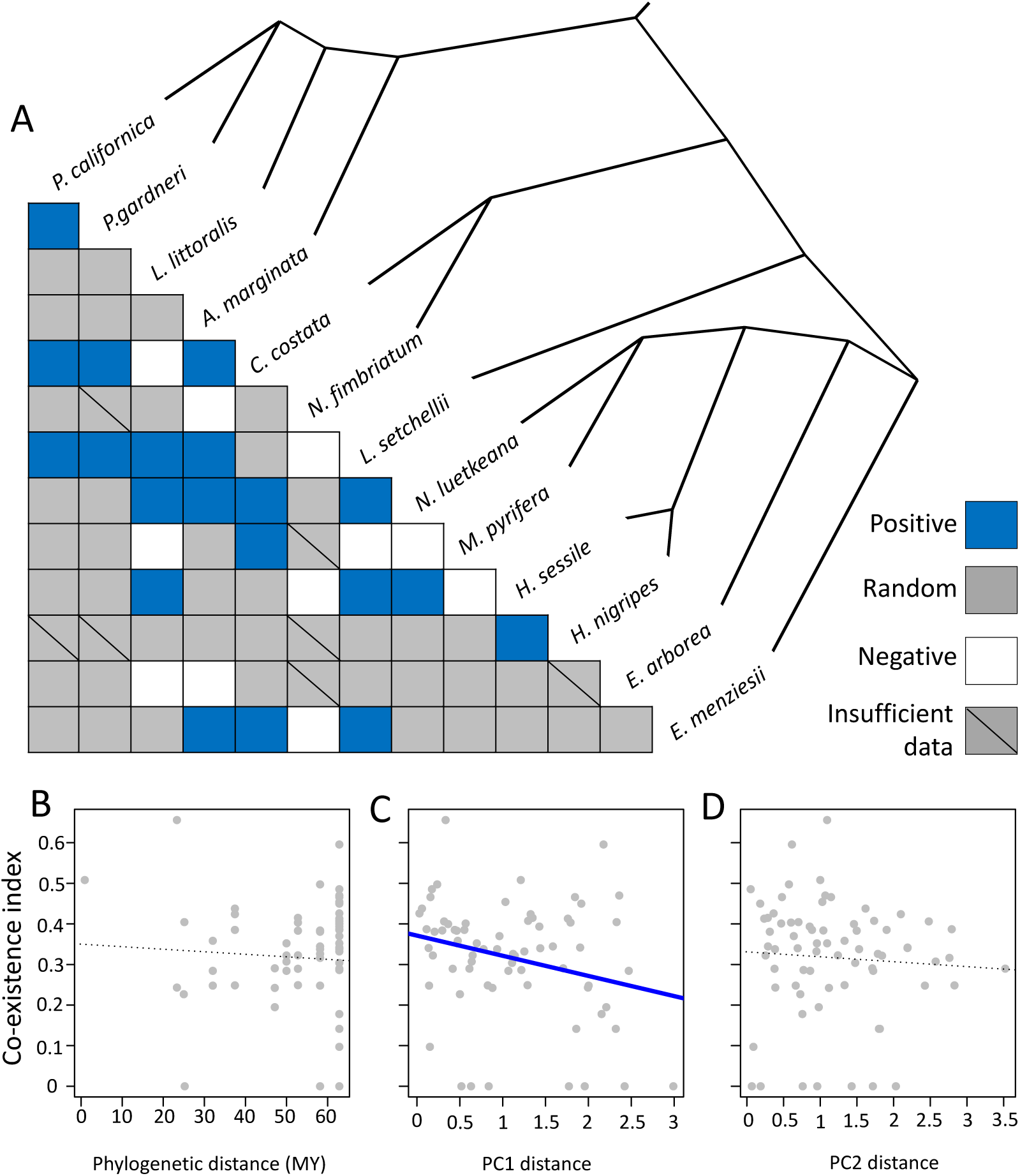
(A) Correlation matrix of species pairs. Colour in each cell indicates whether there was a significant positive or negative correlation between the occurrences of each pair of species, after correcting for false detection rate (q < 0.05). (B-D) Co-occurrence index [Log ((observed co-occurrence / expected co-occurrence) + 1)] versus (A) phylogenetic distance between species pairs in millions of years, (B) distance in PC1 for each species pair and, (C) distance in PC2 for each species pair. Dotted lines indicate insignificant trends, while the solid blue line in panel B indicates a significant slope (P < 0.05).

Despite clear evidence of non-random community assembly, there was no effect of phylogenetic distance on the probability of co-occurrence between species. The only significant predictor of pairwise non-random co-occurrence (measured as “co-occurrence index”) was distance in PC1 between species pairs (Linear regression: F=5.075, df=69, P=0.02746; Fig 3C). Phylogenetic distance (Linear regression: F=0.2392, df=69, P=0.6263; Fig 3B) and PC2 distances (Linear regression: F=0.3037, df=69, P=0.5833; Fig 3D) did not significantly correlate with the pairwise co-occurrence of species.

There was a significant relationship between average wave exposure of a species and its value of PC1 (Linear model: F = 6.809, df = 1 and 12, P = 0.0228; PGLS model: t = 3.9823, df = 14 and 2, P = 0.002; Fig 4), but not PC2 (Linear model: F = 0.1225, df=1 and 12, P = 0.732; PGLS model: t = 0.8316, df = 14 and 2, P = 0.4219), such that structurally reinforced species tended to be found at more wave exposed sites. This relationship was significant even when removing *Postelsia palmaeformis*, the strongest and most wave tolerant species, from the analysis (Linear model: F = 5.161, df = 1 and 11, P = 0.0441; PGLS model: t = 3.0250, df = 13 and 2, P = 0.0116). The only traits that significantly correlated with the average wave exposure of a species on their own were HMF and ε (Table 2). There was a possible, but not significant negative correlation between blade stiffness and average wave exposure.

**Fig 4.**
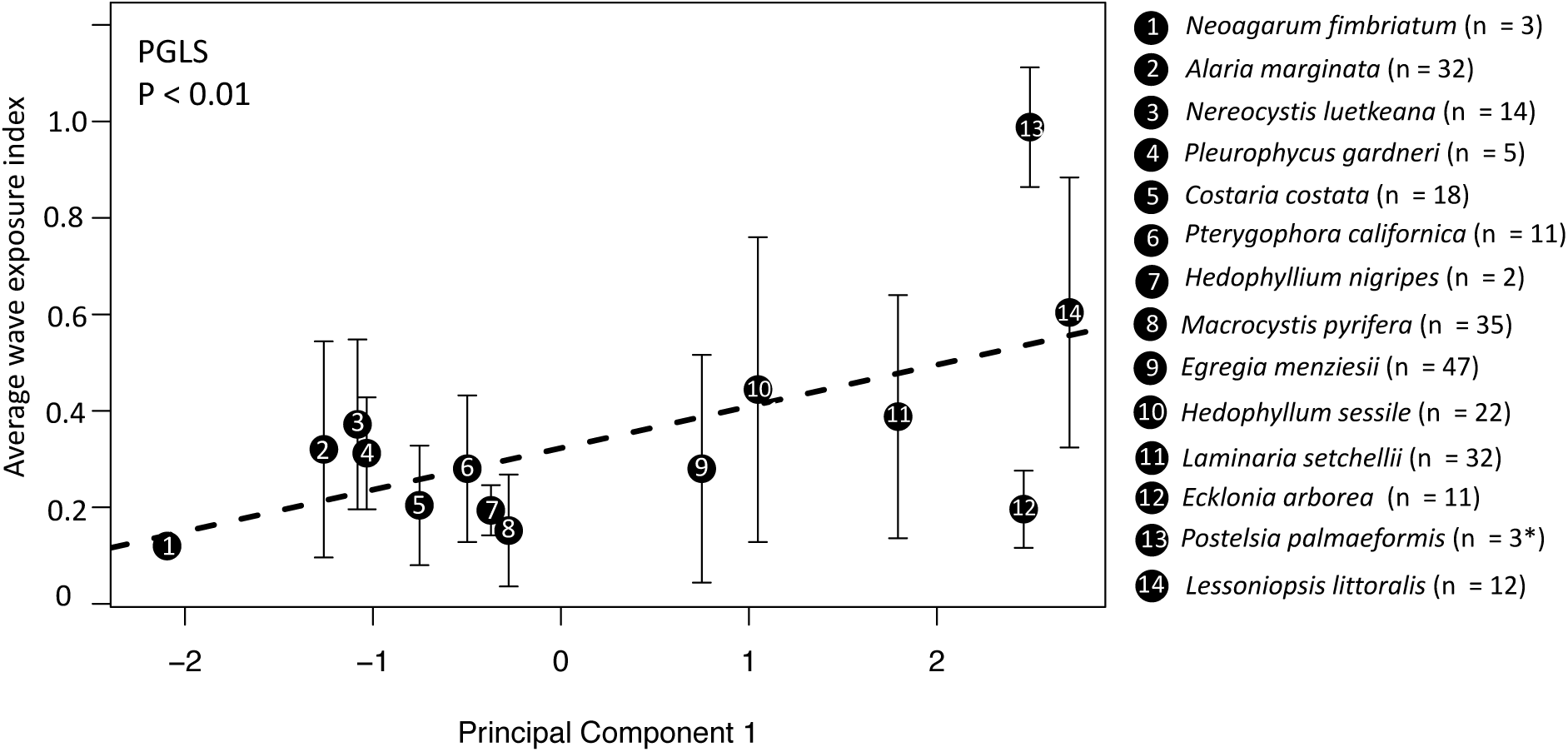
Relationship between wave exposure and principal component 1. Data points represent the average wave exposure that a species was found at (+/− variance) plotted against its value of PC1.

**Table 2.**
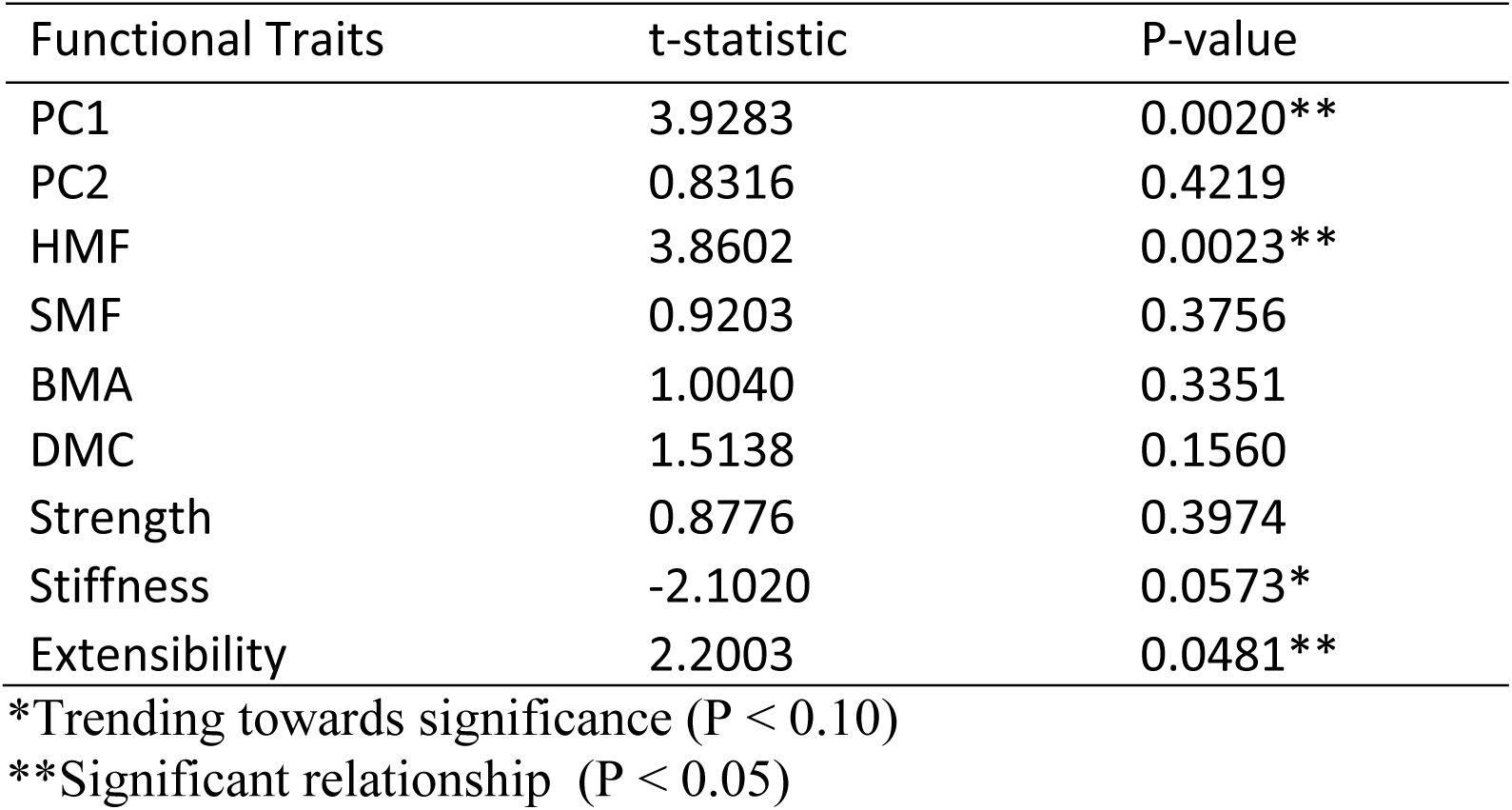
Results of PGLS models testing for correlations between traits and average wave exposure of species (df = 1,12)

### Kelp species are phylogenetically overdispersed across local communities

Use of phylogenetic indices demonstrate that no communities examined were significantly phylogenetically clustered and most communities trended towards phylogenetic overdispersion relative to simulations (Fig 5). Although only a few sites were significantly overdispersed (NRI: n = 3, NTI = 7; Fig 5), average phylogenetic NRI and NTI values were significantly different from zero (NRI: t-test: t = 3.917, df = 86, p = 0.00018; NTI: t-test: t = 9.4708, df = 86, p < 0.0001). The few communities that trended towards phylogenetic clustering were composed of only a small number of species, where clustering of species at sites appeared to be random on average (NTI and NRI approximate zero).

**Fig 5.**
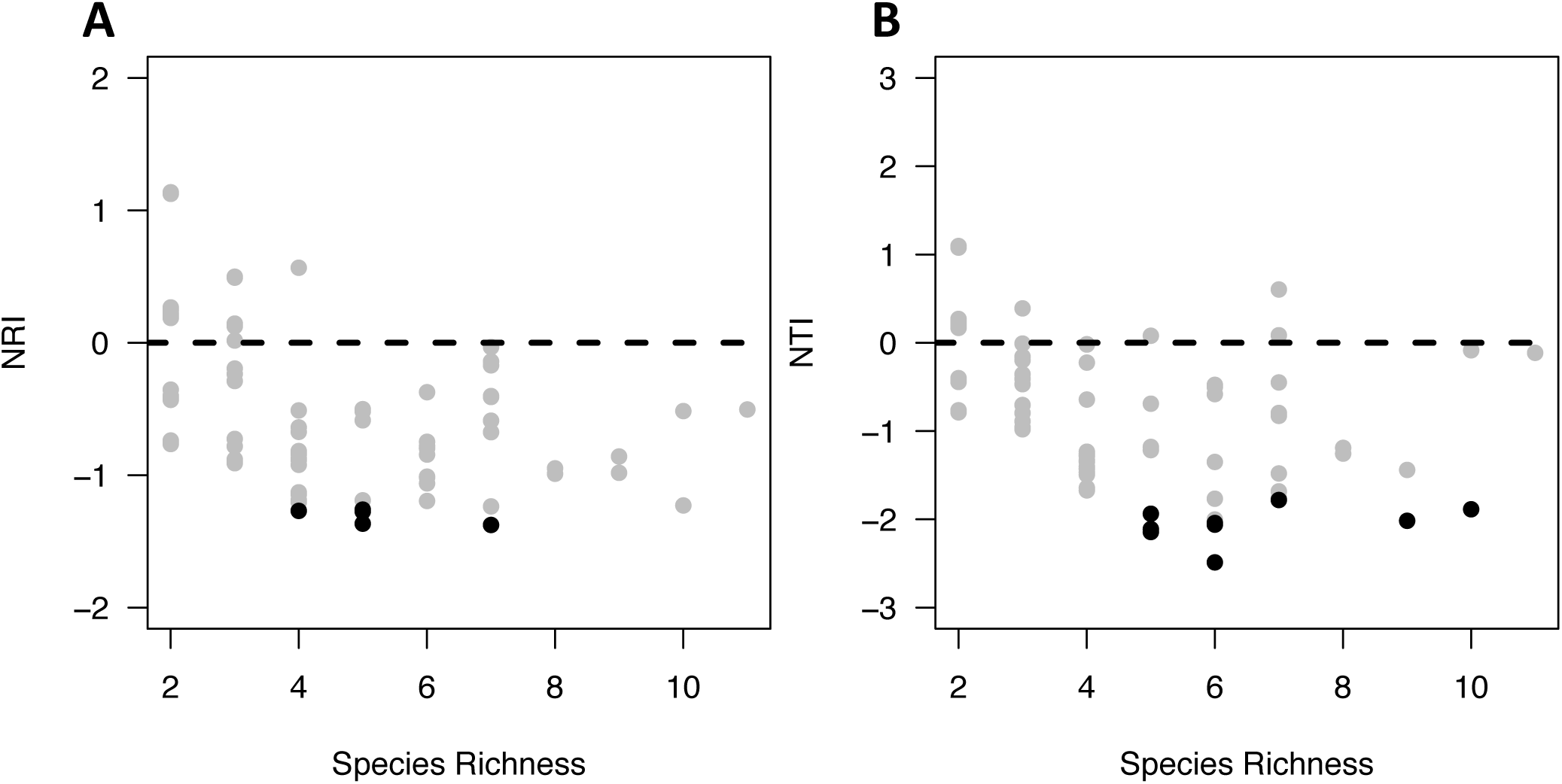
Metrics of phylogenetic community assembly (NRI and NTI) plotted against the species richness of communities. Data points represent individual communities and significance is indicated with dot colour. Black dots indicate that communities are significantly structured by phylogeny, while grey dots indicate no significant phylogenetic effect.

Binomial models of species presence and absence along a continuous wave exposure axis further demonstrates how species in each subclade have convergently adapted to different regimes of wave exposure (Fig. 6). Individual species clearly varied in distribution across the gradient of wave exposure and closely related species (e.g. *Nereocystis luetkeana* and *Macrocystis pyrifera*) tended to specialize in different wave exposure regimes. The clear exception here is the species pair *Pterygophora californica* and *Pleurophycus gardneri* that are sisters and had nearly identical distributions across the wave exposure gradient (Fig 6). There was a significant effect of wave exposure category on community composition (PERMANOVA: F =13.205, P < 0.001; Fig S5), indicating that differences in species distributions across the wave exposure gradient scale up to community level differences in species composition at wave exposed versus wave sheltered sites.

**Fig 6.**
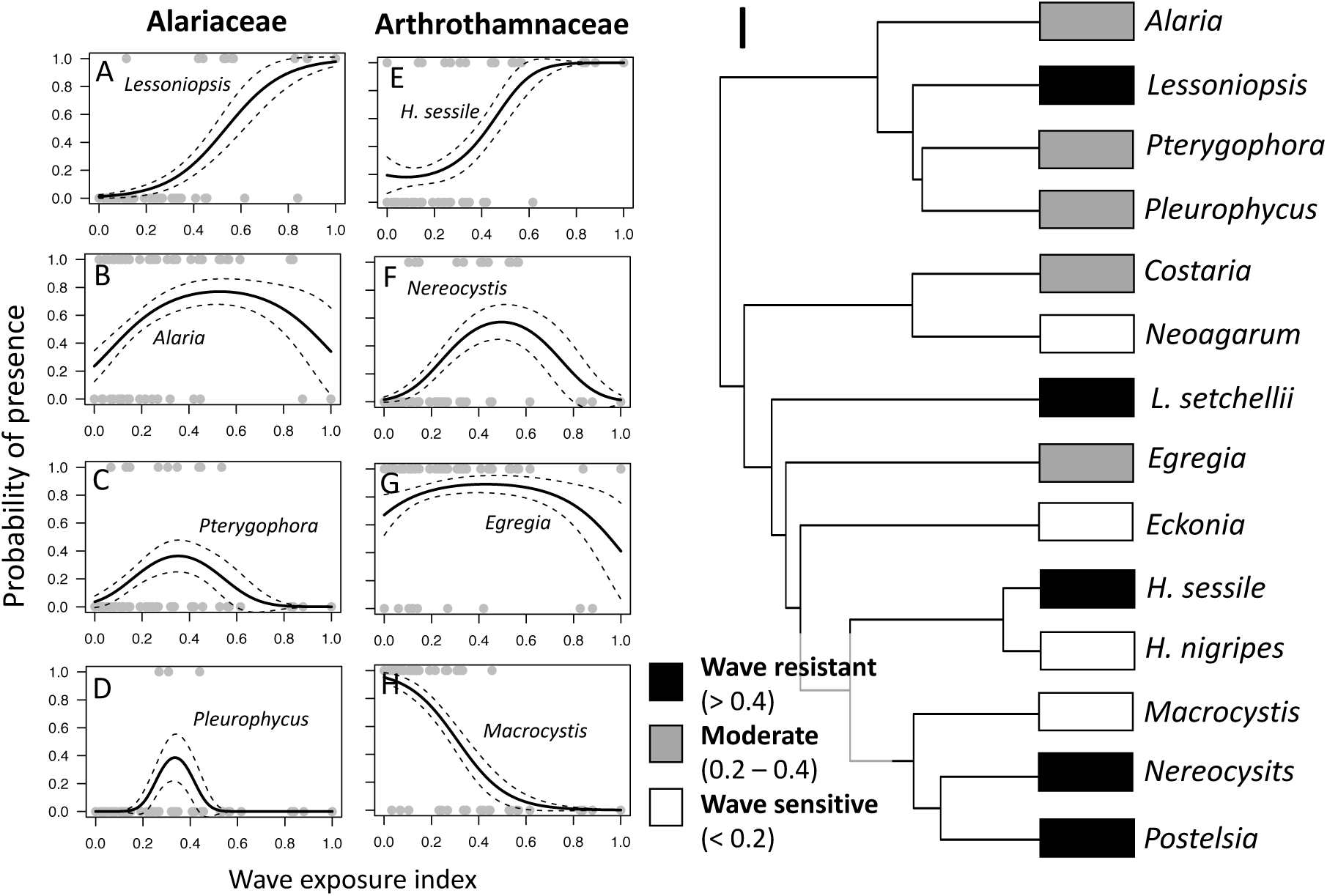
(A-H) Logistic polynomial regressions of species occupancy across a gradient of wave exposure. Columns represent members of two different kelp families (left = Alariaceae, right = Arthrothamnaceae). (I) Phylogeny of the kelps with average wave exposure split into three categories: wave resistant species, moderate species and wave sensitive species.

## Discussion

We demonstrated evolutionary lability in the traits underlying the habitat niches of kelps and suggest that this has resulted in convergent patterns of habitat use across species. Neither principal component, nor any of the individual traits that make them up, were found to be phylogenetically conserved across species (Table 1). In fact, for all traits, Blomberg’s K was less than 1, suggesting that closely related species are more dissimilar than predicted by the null model (although not significant). Yet, PC1 (structural reinforcement) was a predictor of both pairwise species co-occurrences (Fig 3) and the position of individual species along the gradient of wave exposure (Fig 4), indicating a role of structural traits in determining the habitat niche of species. We further propose that this relationship is causal on the basis that many field studies and biomechanical models have demonstrated the role of rapid water motion as a strong selective pressure for increased tolerance to physical forces (Johnson and Koehl 1994, Duggins et al. 2003, Martone et al. 2012, Demes et al. 2013, Starko et al. 2014).

Across communities, species co-occurrence patterns reflect the influence of environmental filtering on community composition but result in overdispersion, rather than clustering, of closely related species. Phylogenetic community indices (NRI and NTI) reveal that communities are made up of more distantly related species than predicted (Fig 5), indicative of phylogenetic overdispersion across kelp communities. Phylogenetic overdispersion of communities is commonly interpreted as phenotypic overdispersion and treated as evidence for competitive exclusion (e.g. Webb 2000, Webb et al. 2002, Cooper et al. 2008). The idea here being that species with similar niches will be unable to co-exist if competition is an important driver of community assembly. However, in our study, species that were commonly found together also tended to be those with correlated niches, indicative of environmental filtering, not competition. For example, *Lessoniopsis littoralis* and *Saccharina sessilis*, two distantly related species that have similar values of PC1 and specialize in wave-swept environments (Fig 6), were positively correlated across the community matrix (Fig 3). Conversely, species that specialize in different wave exposure regimes tended to be negatively correlated. For example, *Neoagarum fimbriatum*, a specialist in wave sheltered areas, and *Laminaria setchellii*, a wave exposed specialist (Fig 6), co-occurred significantly less often than predicted (Fig 3). Thus, kelp communities are filtered strongly but the phenotypes that allow species to pass this filter have evolved convergently in different subclades, resulting in communities of species that have similar phenotypes but come from different clades.

Past work has suggested that traits associated with habitat niche are highly conserved while α niche traits, which result in co-existence of taxa, are more labile (Silvertown et al. 2006a, 2006b; but see Cavender-Bares et al. 2004a). While this framework may hold in many groups of embryophytes, we show that this is not the case for kelps. Habitat niche traits in the kelps are labile and tend to be largely dissimilar among close relatives (Fig 6). While the basis of convergence in traits can be challenging to interpret and may differ across taxa, we propose that partitioning of habitats is an important means by which kelps achieve reproductive isolation and undergo speciation. Partitioning can occur either through character displacement, where competition between close relatives drives the weaker competitor to adapt to new environments (Brown and Wilson 1956), or through the splitting of a generalist niche into multiple specialized niches (Funk 1998). There is substantial evidence that simultaneous phenotypic and genetic divergence across the kelps is common and may be an important driver of diversification. In Table 3, we describe five known instances where partitioning along a gradient of wave exposure has resulted in genetic differentiation of populations or incipient speciation. The prevalence of this pattern in ongoing or incipient speciation events, lends support to our hypothesis that niche partitioning along a wave exposure gradients has been a repeated driver of sympatric speciation and that these processes observed in past studies near the tips of the phylogeny scale up to explain patterns of niche evolution across the broader kelp phylogeny. Close relatives may specialize in different positions along environmental gradients, leading to parallel adaptive radiation across subclades, possibly helping to maintain coexistence of species across broad geographic scales (MacArthur 1958, Cavender-Bares et al. 2004b, 2004a, Losos 2008, Cavender-Bares et al. 2018). We further hypothesize that α niche traits may be more conserved than β niche traits across the kelps, leading to increased co-existence between distant relatives. While it is unclear exactly what traits would promote co-existence across kelp species, morphological features such as the presence of buoyant floats or long, rigid stipes may be somewhat more conserved than the traits examined here, despite multiple origins (Starko et al 2019b). Differences in stature within the water column have been linked to competitive hierarchies in kelps (Edwards and Connell 2015) and may thus make up a component of species α niches.

**Table 3.** Evidence of incipient speciation occurring across gradients of wave exposure

| Species | Environmental gradient | Description | Evidence of differentiation | References |
| --- | --- | --- | --- | --- |
| *Ecklonia arborea* | Wave exposure | Genetic differentiation associated with changes in blade morphology and wave exposure | M13 DNA Fingerprinting | Roberson & Coyer 2004 |
| *Egregia menziesii* | Wave exposure, latitude | Difference in blade and rachis morphology at wave exposed versus sheltered sites; evidence of differential mortality depending on morphology | No direct evidence of genetic differentiation with ITS, despite parapatric overlap of populations. Reciprocal transplants suggest phenotype is genetically determined | Blanchette et al. 2002, Henkel et al. 2007 |
| *Macrocystis pyrifera* | Wave exposure, outer versus inner coast | Difference between wave exposed and wave sheltered morphs; phenotypic-genetic correlations among juveniles suggest local adaptation and differentiation | Genetic distance in ITS2 and microsatellites; Spatially isolated (outer coast vs. Sea of Chiloe) | Kopczak et al. 1991, Astorga et al. 2012, Camus et al. 2018 |
| *Pelagophycus porra* | Wave exposure, substrate | Two distinct morphologies known from the Channel Islands, one on wave exposed sides of islands, the other from wave protected sides. Exposed sites are rocky, sheltered sites are mixed with soft sediment | Random amplified polymorphic DNA show isolation, ITS shows no differentiation | Miller et al. 2000 |
| *Saccharina latissima* sensu lato | Wave exposure | A wave-exposed specialist population from Maine was described as new species, *Saccharina angustissima*, making *S. latissima* paraphyletic | Difference in rbcL and cox3 (but not cox1) between *S. angustissima* and *S. latissima* populations from Maine; common garden revealed that blade shape is genetically determined | Augyte et al. 2018 |

Multiple hypotheses may explain why phenotypic divergence, rather than niche conservatism, is the dominant process behind kelp phenotypic evolution. Kelps diversified only recently and following massive changes to global climate (Starko et al. 2019b). Kelps are much larger and more competitive than other macroalgal species (Edwards and Connell 2015) but rely on cool waters and an abundance of nutrients. Cooling of the oceans may have created an ecological opportunity for kelps, allowing them to diversify across and dominate rocky shores throughout the Northeast Pacific (Bolton 2010, Starko et al. 2019b, Vermeij et al. 2019). This ecological opportunity may have promoted selection for niche partitioning as has been documented previously, such as in oak trees (Cavender-Bares et al. 2004a, Cavender-Bares et al. 2018), the silversword alliance (Ackerly 2009, Blonder et al. 2016) and Carribean anoles (Losos et al. 2003). If this is the case, then it is because of (and not in spite of) the ecological relevance of these traits that we find no phylogenetic signal. This hypothesis is further supported by recent evidence that temperature tolerance and chemical deterrent production, which determine the geographic range limits of species and the responses of species to herbivory, respectively, are also highly labile across kelps (heat tolerance: Muth et al. 2019, chemical deterrents: Starko et al. 2019b). An alternative hypothesis is that these patterns are typical of marine macroalgae that to date have been poorly explored in this regard. Individual macroalgae are fixed in place but lineages can span broad gradients of stress and disturbance, relying only on relatively simple morphological adaptations to survive. Because traits are generally simple, novelty may not be particularly important in determining the habitat niche of macroalgae, and thus strong selection on quantitative, heritable traits may lead to divergence being common among close relatives. This hypothesis is supported by recent work on coralline algae, showing that intense grazing by urchins (analogous to environmental filtering) does not lead to phylogenetic clustering (Hind et al. 2019) as predicted by assumptions of niche conservatism. Regardless of the generality of our results to other marine macroalgae, we show that niche partitioning has been an important driver of kelp phenotypic evolution, highlighting the importance of divergent selection in the evolution of a lineage of marine foundation species. Future work should investigate the extent to which these patterns extent to other marine lineages in order to determine how ecological and evolutionary processes interact in the ocean.

### Conclusions

We demonstrate that the distribution of phenotypic traits across the kelp phylogeny represents convergent evolution of niche structure. We propose that this is a consequence of niche partitioning by close relatives, with wave exposure as an important axis of niche structure. More broadly, our results provide clear evidence that traits are not always phylogenetically conserved and that phylogenies are not proxies for ecological differences between species, but instead provide an opportunity to explore how local scale processes influence macroevolutionary diversification (as argued by Gerhold et al. 2015). Phenotypic divergence between close relatives may be expected in particular situations and therefore understanding the circumstances and spatial scales at which phenotypic conservatism or divergence are expected is the critical next step for the field of phylogenetic community ecology.

## Acknowledgements

The authors would like to thank L.Campbell, L.Bailey, E.Creviston, K.James, A.Warren, M.Brophy, A.Danasel, M.Fass, J.Townsend, E.Hardy, E.Sadler, L.Gonzalez, J.Matsushiba, L. Liggan, R. Munger, and B. Radziej for help in the field, L.Gendall for assistance with visualization and E. Clelland, S. Gray and the rest of the BMSC staff for assistance with field logistics. This work would not have been possible without the field surveys conducted by L. Druehl and C. Elliot, with funding from Parks Canada. Funding for this project was provided by the Natural Sciences and Engineering Research Council (NSERC) in the form of a Discovery Grant (to PTM), and a Canadian Postgraduate Scholarship (to SS), as well as from the Killam Trust in the form of an Izaak Killam Pre-doctoral Fellowship (to SS). The use of laboratory equipment was also supported by the Canadian Foundation for Innovation in the form of a Leaders Opportunity Fund Grant (#27431) to PTM. Research conducted in Pacific Rim National Park was done with permission from Parks Canada (Permit Numbers: PRN-2015-18843, PRN-2018-29480). Thanks also to Huu-ay-aht First Nations for allowing research on their lands. We thank Q. Cronk, S. Graham, C. Harley and H. Verbruggen for helpful feedback on an earlier version of this manuscript.

## Supplemental Information

**Table S1.** Locations of field sites from which trait data were collected on different species

| Site Name | Location | Latitude | Longitude |
| --- | --- | --- | --- |
| Bamfield Inlet | Barkley Sound, BC | 48.8345 | -125.13682 |
| Brady's Blowhole | Barkley Sound, BC | 48.82329 | -125.16151 |
| Edward King Island | Barkley Sound, BC | 48.82235 | -125.21731 |
| Scott's Bay | Barkley Sound, BC | 48.83413 | -125.14775 |
| Prasiola Point | Barkley Sound, BC | 48.81751 | -125.16926 |
| Cape Beale | Barkley Sound, BC | 48.78537 | -125.2165 |
| Ogden Point | Victoria, BC | 48.41399 | -123.38572 |
| Botanical Beach | Port Renfrew, BC | 48.52753 | -124.44877 |
| Whytecliff Park | Vancouver, BC | 49.37226 | -123.29212 |

**Table S2.** Sources of trait data used in this study. σ = breaking stress, *E* = tensile modulus (stiffness), ε = extensibility, SMF = stipe mass fraction, HMF = holdfast mass fraction, DMC = dry matter content of blades, BMA = blade mass per area.

| Species | Materials ($\sigma$, $E$ & $\epsilon$) | Biomass (SMF & HMF) | Blade Properties (DMC & LMA) |
| --- | --- | --- | --- |
| *Alaria marginata* | This study, Botanical Beach (n = 8) | Starko & Martone 2016 (n = 5) | This study, Blowhole (n = 10) |
| *Lessoniopsis littoralis* | This study, Brady's Blowhole (n = 5) | Starko & Martone 2016 (n = 5) | This study, Blowhole (n = 9) |
| *Pleurophycus gardneri* | This study, Ogden Point (n = 5) | Starko & Martone 2016 (n = 5) | This study, Ogden Point (n = 6) |
| *Pterygophora californica* | This study, Botanical Beach (n = 8) | Starko & Martone 2016 (n = 5) | This study, Ogden Point (n = 2) |
| *Costaria costata* | This study, Whytecliff Park (n = 8) | Starko & Martone 2016 (n = 5) | This study, Scott's Bay (n = 3) |
| *Neoagarum fimbriatum* | This study, Whytecliff Park (n = 8) | Starko & Martone 2016 (n = 5) | This study, Bamfield Inlet (n = 4) |
| *Egregia menziesii* | Demes et al 2013 (n = 39) | Starko & Martone 2016 (n = 5) | This study, Scott's Bay (n = 5) |
| *Ecklonia arborea* | Hale 2001 | This study, Scott's Bay (n = 5) | This study, Scott's Bay (n = 5) |
| *Cymathaere triplicata* | This study, Ogden Point (n = 7) | This study, Ogden Point (n = 5) | This study, Ogden Point (n = 5) |
| *Nereocystis luetkeana* | This study, Botanical Beach (n = 8) | Starko & Martone 2016 (n = 5) | This study, Scott's Bay (n = 3) |
| *Macrocystis pyrifera* | Hale 2001 | Starko & Martone 2016 (n = 5) | This study, Scott's Bay (n = 13) |
| *Postelsia palmaeformis* | This study, Botanical Beach (n = 8) | Starko & Martone 2016 (n = 5) | This study, Cape Beale (n = 3) |
| *Saccharina sessilis* | This study, Botanical Beach (n = 8) | Starko & Martone 2016 (n = 5) | This study, Prasiola Point (n = 7) |
| *Saccharina nigripes* | This study, Scott's Bay (n = 6) | Starko & Martone 2016 (n = 5) | This study, Scott's Bay (n = 12) |
| *Saccharina latissima* | This study, Bamfield Inlet (n = 4) | Starko & Martone 2016 (n = 4) | This study, Bamfield Inlet (n = 6) |
| *Laminaria setchellii* | Starko et al 2018 (Blowhole) (n = 6) | Starko & Martone 2016 (n = 5) | This study, Blowhole (n = 3) |
| *Laminaria ephemera* | This study ((n = 4) | Starko & Martone 2016 (n = 5) | This study, Edward King (n = 4) |

**Fig S1.**
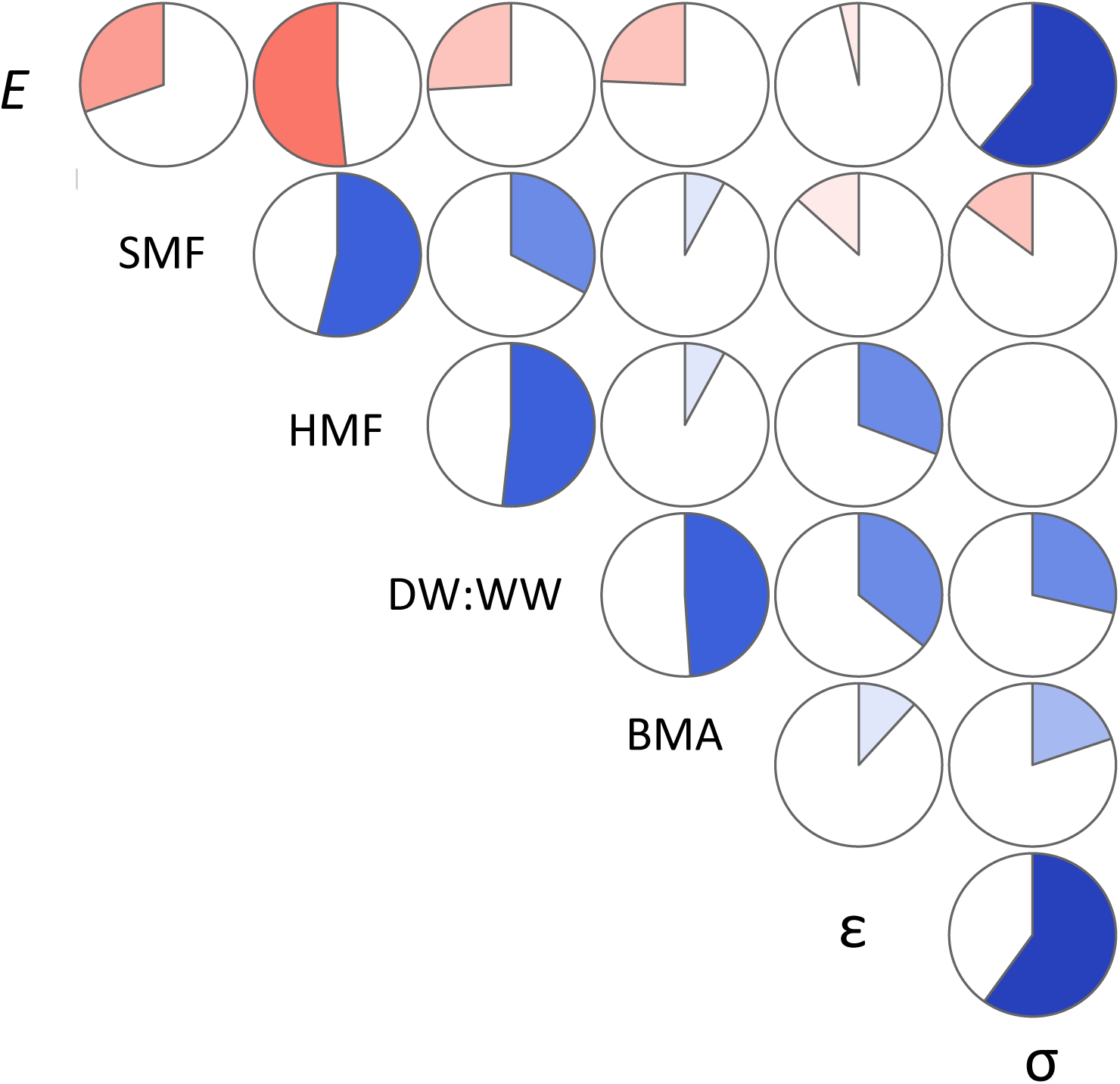
Correlogram of the seven functional traits examined in this study. The filled in pie slices indicate the correlation coefficient, *r* (0 < *r* < 1). Blue slices indicate a positive correlation between traits, while red slices indicate negative correlations.

**Fig S2.**
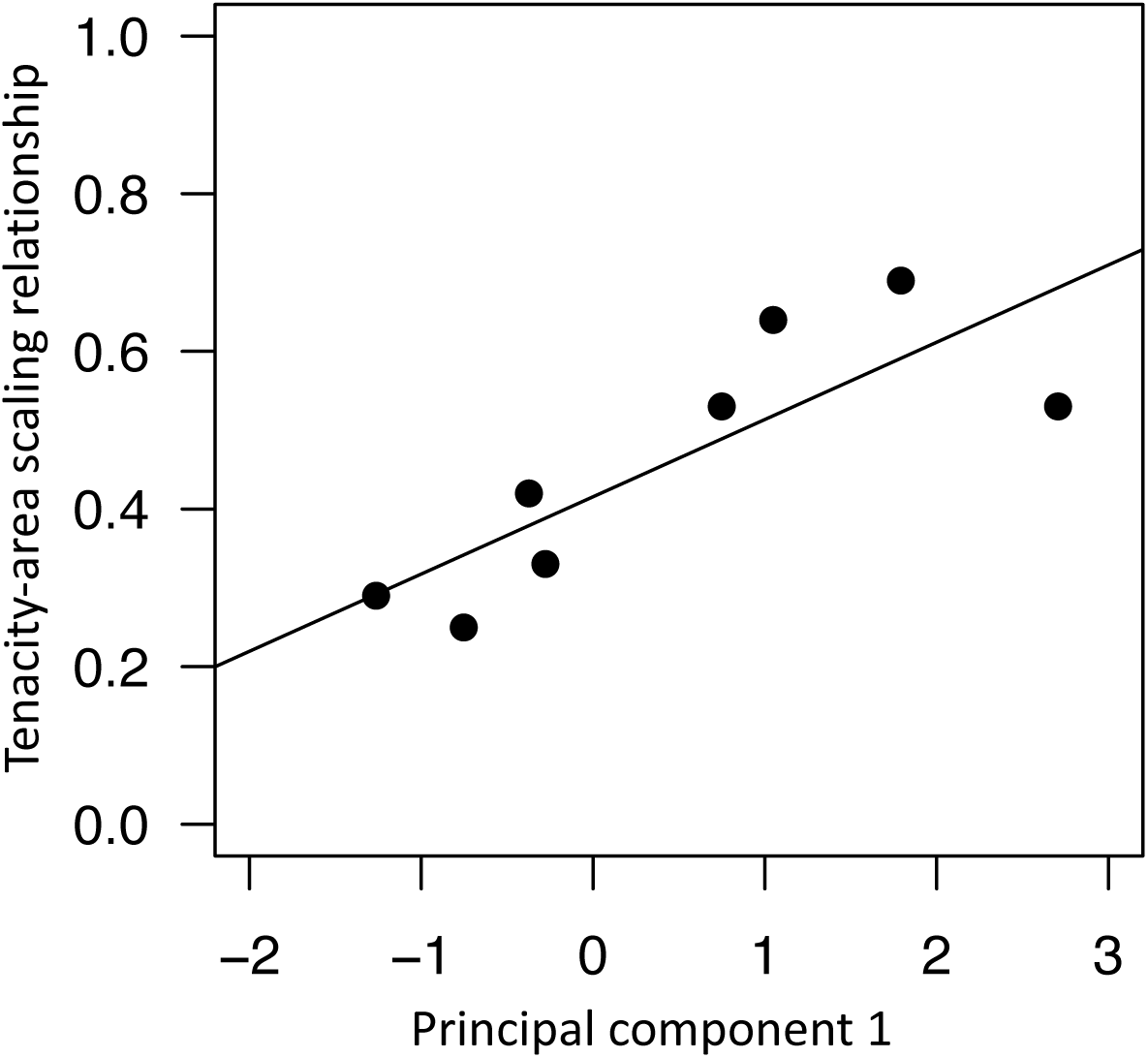
Correlation between principal component 1 (from this study) and tenacity-area scaling relationships (from Starko & Martone 2016).

**Fig S3.**
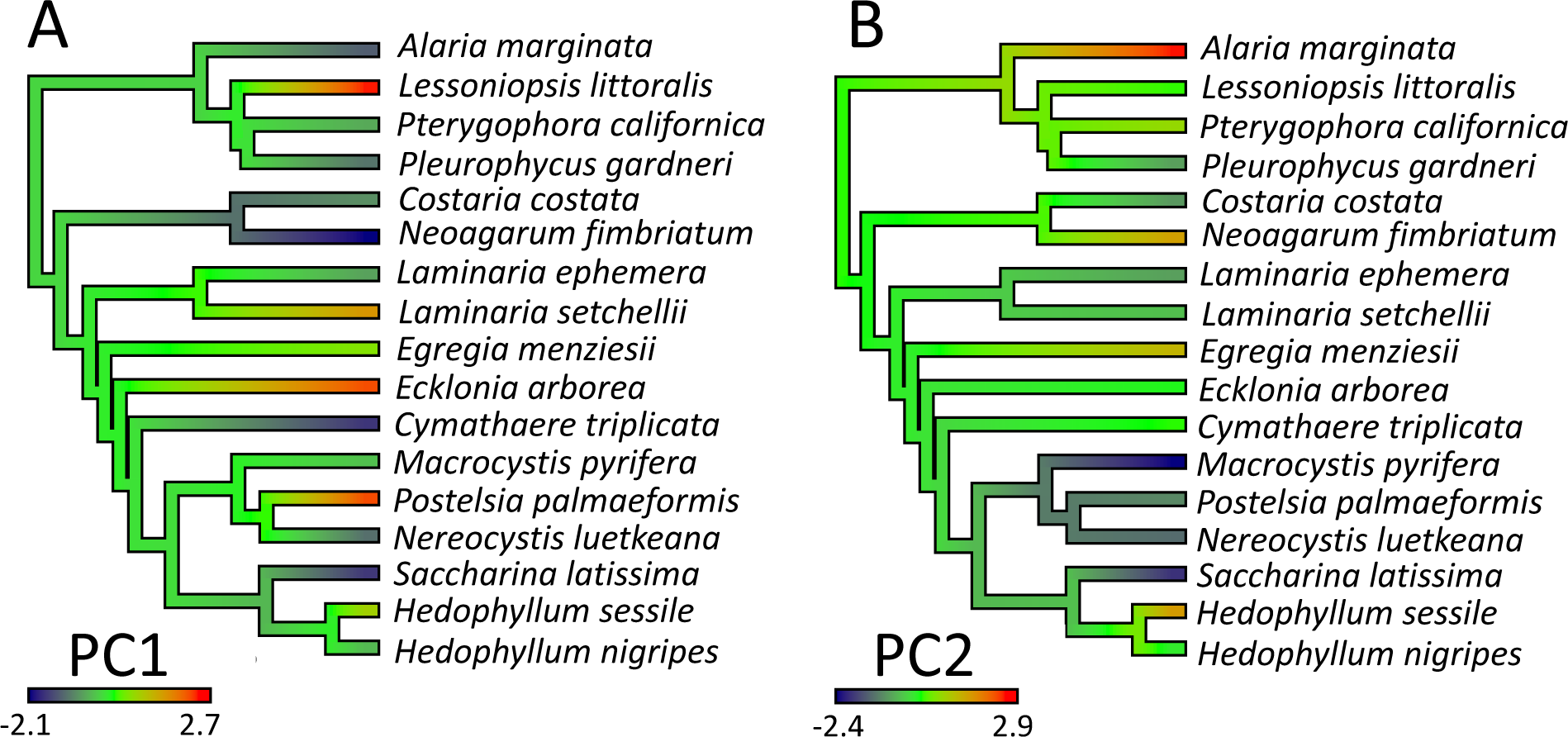
Ancestral state reconstruction of principal components (PC1 and PC2) computed in StableTraits and visualized using contMap. PC1 represents structural reinforcement of the whole kelp thallus, while PC2 represents a component of material properties of the blade.

**Fig S4.**
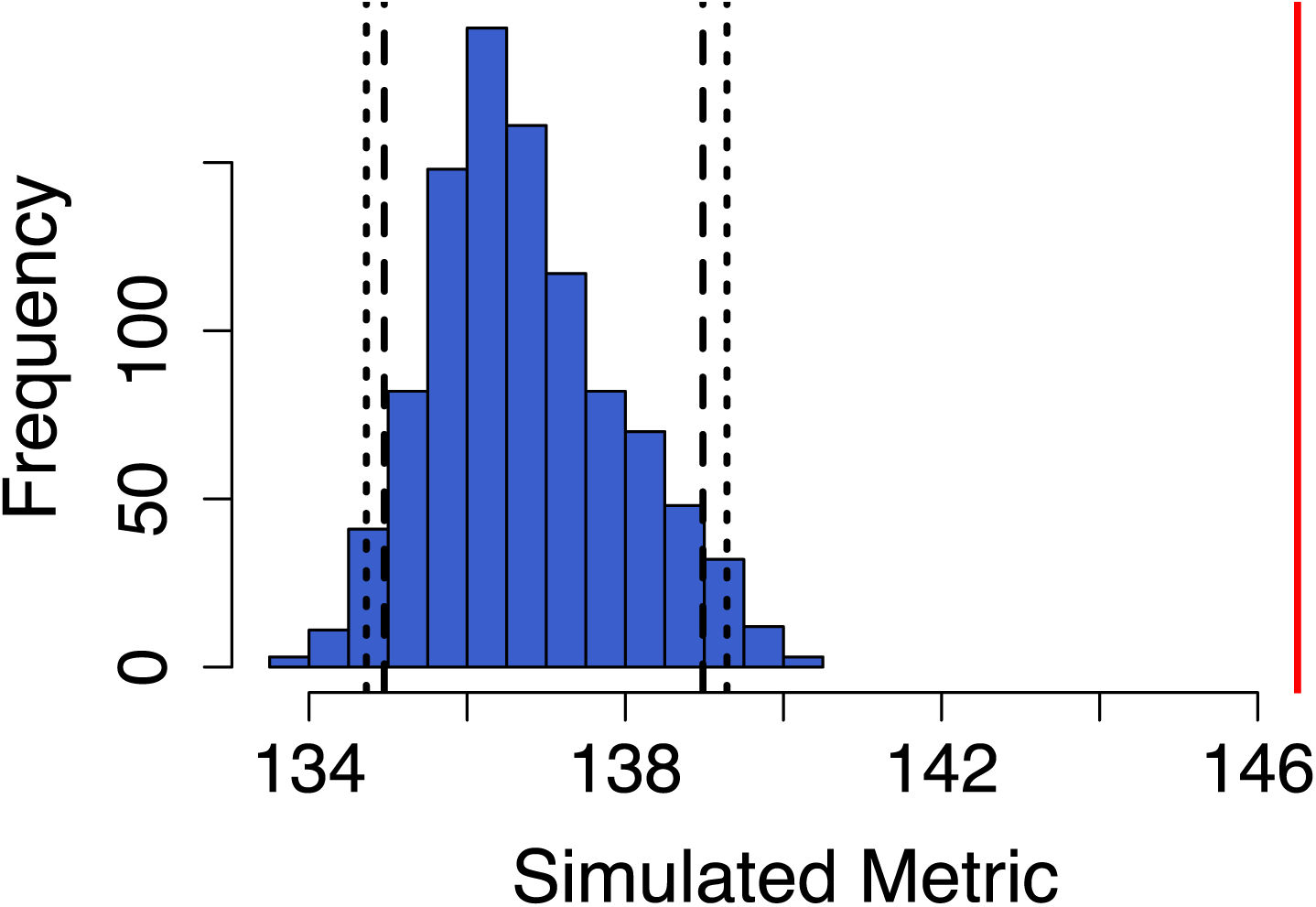
Observed c-score of the community matrix (red line) compared to simulated c-score values from 1000 random communities (blue histogram). Dotted lines indicate 95% confidence intervals. Analysis was conducted in the package EcoSimR.

**Fig S5.**
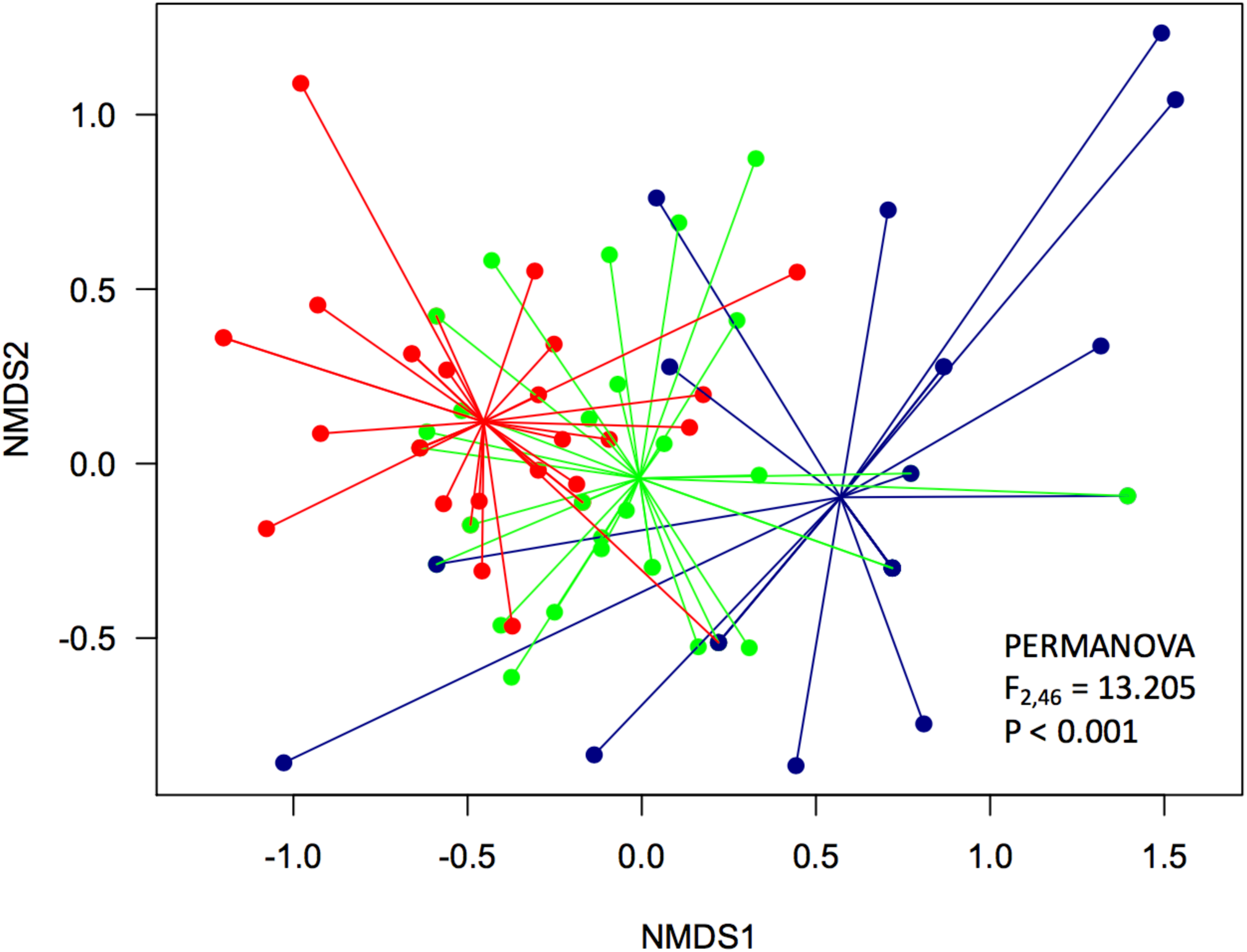
NMDS plot of community presence data for kelp species at 87 sites in Barkley Sound, British Columbia. Sites are coloured by wave exposure category (red = exposed, green = moderate, blue = sheltered) and lines are drawn between all sites and the centroid of its wave exposure category.

